# Episodic pain in Fabry disease is mediated by a heat shock protein-TRPA1 axis

**DOI:** 10.1101/2025.02.20.639340

**Authors:** Jonathan D Enders, Eve K Prodoehl, Signe M Penn, Anvitha Sriram, Cheryl L Stucky

## Abstract

Two-thirds of patients with Fabry disease suffer debilitating pain attacks triggered by exercise, fever, and exposure to environmental heat. These patients face an even greater risk of heat-related episodic pain in the face of global climate change. Almost nothing is known about the biological mechanisms underlying heat-induced pain crises in Fabry disease, and there is no preclinical model available to study Fabry crises. Here, we established the first model of heat-induced pain attacks in Fabry disease by exposing transgenic Fabry rats to environmental heat. Heat exposure precipitated robust mechanical hypersensitivity, closely matching temporal features reported by patients with Fabry disease. At the cellular level, heat exposure sensitized Fabry dorsal root ganglia (DRG) neurons to agonists for transient receptor potential cation channel A1 (TRPA1), but not TRPV1. The heat shock response, which normally confers heat-resilience, was impaired in Fabry disease, and we demonstrated that heat shock proteins (HSP70 and HSP90) regulate TRPA1. Strikingly, pharmacologically inhibiting HSP90 completely prevented cellular and behavioral sensitization by environmental heat in Fabry disease. Together, this work establishes the first model of episodic pain in Fabry disease, implicates the heat shock response in heat-evoked pain episodes, and identifies a novel heat shock protein-TRPA1 regulatory axis.

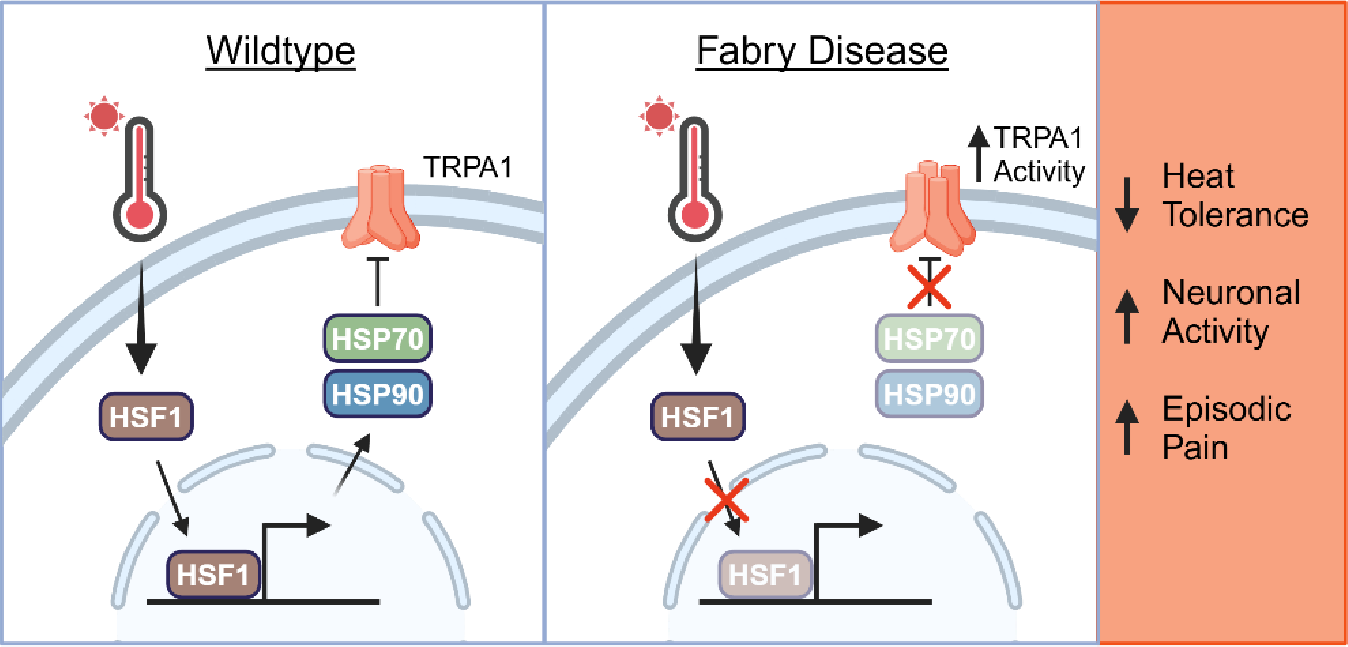

## Introduction

Fabry disease is the most common lysosomal storage disorder, affecting between 1:895-3100(1, 2) live male births. Fabry disease arises from X-linked mutations in the *GLA* gene that reduce α-galactosidase A (α-Gal A) function and subsequently lead to accumulation of its substrate globotriaosylceramide (Gb3). Gb3 accumulates in several tissues, including sensory neurons of the dorsal root ganglia (DRG). Thus, patients with Fabry disease frequently experience chronic pain and episodic pain attacks, also known as Fabry crises. Systematic characterization of pain in Fabry disease revealed that approximately half of adult patients and up to two-thirds of pediatric patients with Fabry disease experience episodic pain(3–5). These pain episodes last between minutes and days(4), severely affect patient lifestyle and quality of life, and are frequently triggered by fever, exercise, and exposure to environmental heat(3–6). There is already a critical need to understand the biological mechanisms that lead to heat-evoked crises in Fabry disease. However, as climate change progresses, patients with Fabry disease risk more frequent exposure to elevated temperatures and, by extension, further increased incidence of heat-induced pain crises. Almost nothing is known about mechanisms underlying Fabry crises, partially due to the lack of established models in which to study episodic pain associated with Fabry disease. Thus, a preclinical model is critically needed in order to study the mechanisms of episodic pain in Fabry disease.

One mechanism that may contribute to episodic pain and heat intolerance in Fabry disease is the heat shock response. The heat shock response is a set of molecular pathways that confer resilience to cellular and environmental stress(7, 8). Stress exposure causes heat shock factor 1 (HSF1) translocation to the nucleus, where it transcriptionally upregulates heat shock proteins(9). Heat shock proteins are molecular chaperones that are classified into different families by size (*e.g.*, HSP40, HSP70, HSP90, HSP110, and small heat shock proteins) and aid protein folding, receptor trafficking, signal transduction, and metabolic function(7, 10–13). Heat shock protein expression or function may be diminished in Fabry disease, as patients with Fabry disease often have heat intolerance. HSP70 family members have been shown to regulate trafficking and turnover of transient potential cation channel vanilloid 1 (TRPV1)(14), which transduces heat stimuli, mediates heat hypersensitivity after tissue inflammation, and contributes to neuropathic pain. Though TRPV1 is not sensitized during the chronic phase of pain associated with Fabry disease(15, 16), it is possible that heat stress sensitizes TRPV1 in Fabry disease. The thermal sensitivity of transient receptor potential cation channel ankyrin-containing 1 (TRPA1), which is closely related to and partially regulated by TRPV1(17), remains controversial. However, we previously demonstrated a role for TRPA1 in pain-like behaviors in Fabry disease at steady state(15), and TRPA1 remains an attractive analgesic target for neuropathic and inflammatory pain(18). It is unknown, however, whether TRPA1 is regulated by heat shock proteins, nor is it known whether TRPV1 or TRPA1 contribute to episodic pain in Fabry disease. Answering these questions will not only identify new therapeutic targets for heat-evoked pain crises in Fabry disease, but might also provide insight to the normal function and regulation of these channels that may be leveraged to treat other conditions of chronic and episodic pain.

Here, we developed a model of Fabry pain crises to explore the molecular basis of the debilitating, heat-evoked episodic pain experienced by patients with Fabry disease. To accomplish this, we applied mild heat stress to *Gla*-knockout (Fabry) rats and DRG neurons. We validated this model by assessing the onset of pain behaviors in Fabry rats following heat exposure, confirming that they closely matched the sensory and temporal features of pain attacks in patients with Fabry disease. Thereafter, we used this foundational model of Fabry heat crises to determine a role for the heat shock response in heat-evoked pain crises and identify a novel regulatory mechanism for TRPA1 by heat shock proteins. Thus, we developed for the first time a model of Fabry pain crises and established the heat shock protein-TRPA1 regulatory axis as a viable therapeutic target for Fabry crises.

## Results

### Transient heat exposure elicits mechanical allodynia in Fabry but not wildtype rats

Patients with Fabry disease frequently experience pain crises evoked by environmental heat(4, 19). To examine whether Fabry rats recapitulated this feature of neuropathic pain in Fabry disease, we compared mechanically evoked pain-like behaviors between wildtype, heterozygous, and homozygous Fabry rats following transient heat treatment (30 minutes at 40°C, **Figure 1A**). At 10-14 weeks of age, Fabry animals did not exhibit statistically different mechanical withdrawal thresholds or responses to noxious needle from wildtype littermate controls at baseline (**Figure 1**). Likewise, there were no differences between wildtype, heterozygote, and homozygous Fabry rats for responses to dynamic brush at baseline. Within one hour of transient heat treatment, hemizygous male and both heterozygous and homozygous female Fabry rats exhibited hypersensitivity to blunt, punctate mechanical stimulation (**Figure 1B, 1E**). This hypersensitivity persisted for at least 4 hours and partially recovered by 24 hours post-heat treatment. We observed a gene dose effect in females, where heterozygous Fabry rats developed pronounced mechanical hypersensitivity but recovered faster than homozygous female Fabry rats. There was also a significant increase in needle-evoked attending behavior (*e.g.*, jumping, guarding, paw flutter, or shaking) 1-hour following heat treatment in male and female Fabry animals, which did not fully resolve within 24-hours of heat treatment (**Figure 1C, 1F**). Heterozygous females likewise exhibited increased needle-evoked attending behaviors 1-, 2-, and 4-hours following heat treatment. Unlike homozygous female animals, however, sensitization to pinprick resolved in heterozygous females within 24-hours of heat treatment. Additionally, both male and female Fabry rats and heterozygote females demonstrated transient brush allodynia following heat treatment (**Figure 1D, 1G**). One hour after heat treatment, there was a significant increase in paw withdrawal and attending behavior to dynamic brush in Fabry but not wildtype rats. This increase in attending behavior persisted for 2-hours post-heat treatment but had largely recovered at the 4- and 24-hour timepoints. There were concomitant decreases in null responses to needle and dynamic brush in Fabry rats follow heat treatment, but no differences in the frequency of simple withdrawal behavior to either stimulus (**Supplemental Figure 1**). Thus, *Gla*-knockout Fabry rats recapitulate the heat-evoked pain episodes experienced by patients.

**Figure 1.**
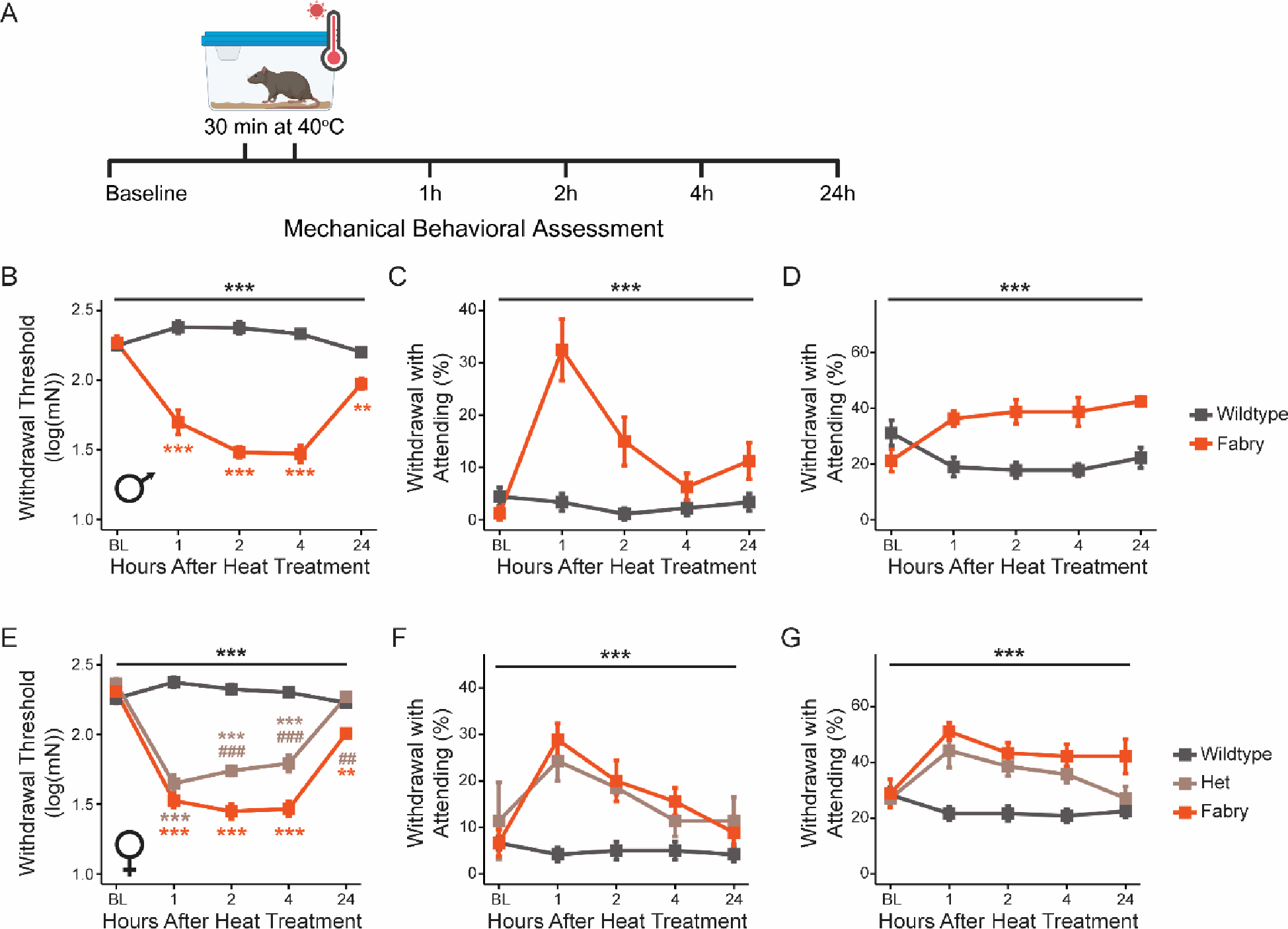
Heat elicits mechanical hypersensitivity in Fabry rats. Mechanical sensitivity was assessed in wildtype and Fabry rats before and after transient heat exposure (40°C for 30 min) (A). Male Fabry rats displayed decreased mechanical withdrawal thresholds (B) and increased attending responses to dynamic brush (C) and needle (D) following heat exposure (n=8-9). Heterozygotes and female Fabry rats also exhibited decreased mechanical withdrawal thresholds (E) and increased attending responses to brush (F) and needle (G). (B-G) 2-Way ANOVA with repeated measure, *** is genotype: p<0.005. (B,E) Tukey’s *post hoc*, colored, ** is comparison to Wildtype: p<0.01, *** is comparison to Wildtype: p<0.005, ## is comparison to Fabry: p<0.01, ### is comparison to Fabry: p<0.005.

### In vitro heat sensitizes TRPA1 in DRG neurons from Fabry rats

We previously reported that TRPA1 is sensitized in Fabry DRG neurons(15), and TRPA1 antagonism rescued steady-state mechanical hypersensitivity in Fabry rats. We therefore asked whether heating isolated DRG neurons further sensitized TRPA1 in Fabry disease (**Figure 2A-D**). Following 30-minute incubation at 40°C, we observed an increased proportion of Fabry, but not wildtype, DRG neurons exhibiting responses to the TRPA1 agonist allyl isothiocyanate (AITC) with calcium transients (**Figure 2C**). The magnitude of AITC-evoked calcium responses was also elevated in Fabry DRG neurons but not wildtype neurons, after heat exposure (**Figure 2D**).

**Figure 2.**
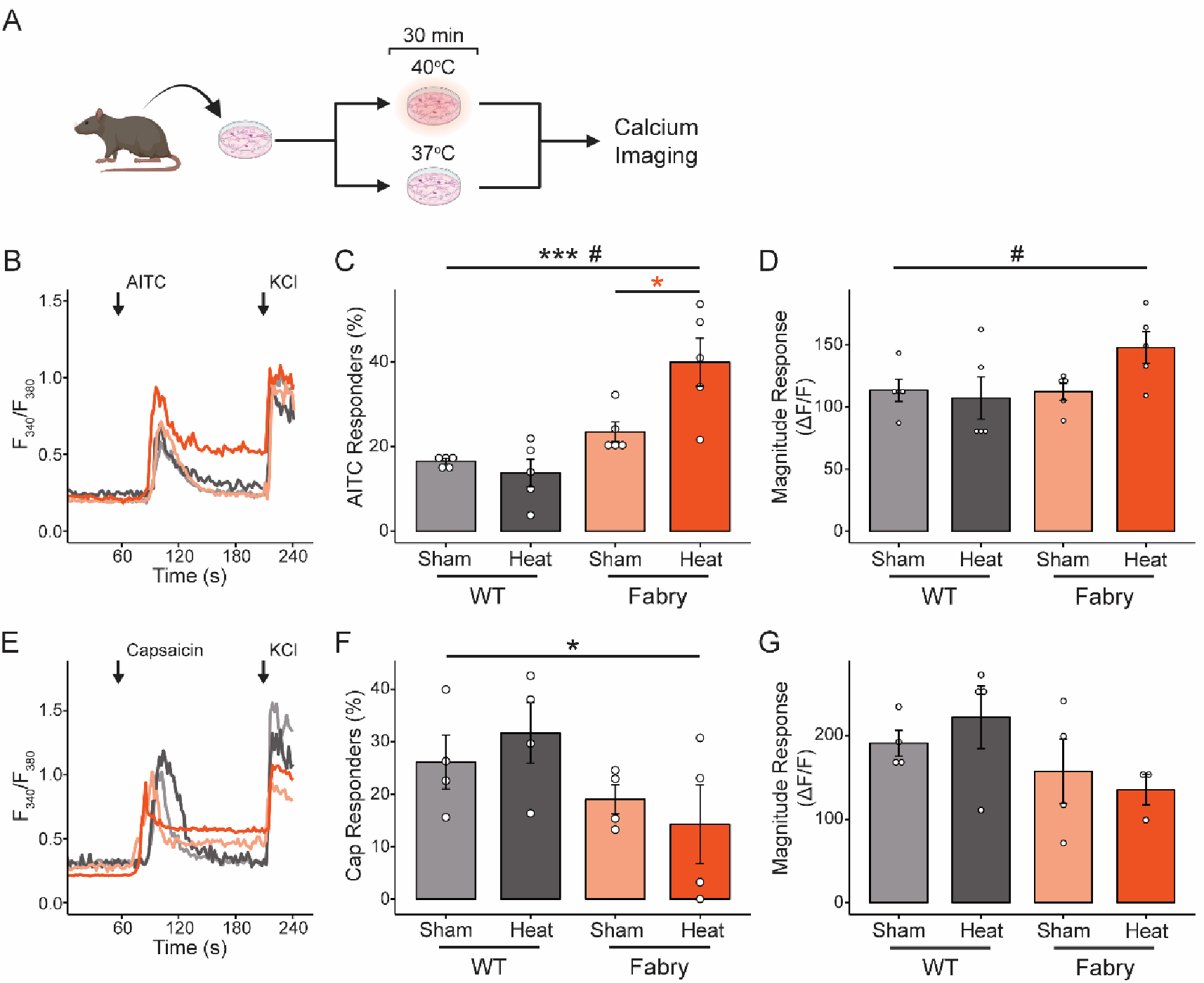
Heat stress sensitizes TRPA1 in Fabry dorsal root ganglia (DRG) neurons. DRG neurons cultured from wildtype or Fabry rats were incubated at 37°C (sham) or 40°C (heat) for 30 minutes one day after plating before calcium imaging studies (A). Representative traces are shown of responses to (B) allyl isothiocyanate (AITC, 30μM) and (E) capsaicin (Cap, 50nM) in DRG neurons. Heat increased the proportion of Fabry DRG neurons responding to AITC (C) and AITC-induced calcium transient magnitude (D). Fabry DRG were less responsive to capsaicin (F) but not different in capsaicin-induced calcium transient magnitude (G) regardless of heat. Each dot represents cultured DRG neurons per rat (n=4-5 rats, 50-90 neurons per rat). 2-way ANOVA with repeated measures, * (black) is genotype: p<0.05, *** (black) is genotype: p<0.005, # is genotype-heat interaction: p<0.05; Tukey *post hoc*, * (orange) is p<0.05.

TRPA1 directly interacts with and is regulated by TRPV1(17). While prior work failed to detect sensitization of TRPV1 in Fabry DRG neurons at steady state(15, 16), TRPV1 is activated near the temperature of heat stress used in these studies(20, 21) and the possibility remains that dysregulation of TRPV1 by heat contributes to TRPA1 sensitization in Fabry disease. Although there were fewer Fabry DRG neurons that responded to the TRPV1 agonist capsaicin with calcium transients, this effect was independent of heat exposure or an interaction between heat and genotype (**Figure 2F**) and there were no changes in the magnitude of capsaicin-induced calcium transients between Fabry and wildtype DRG neurons regardless of heat exposure (**Figure 2G**). We also assessed TRPV4 function in Fabry DRG neurons at steady state and following heat exposure, as TRPV4 detects both warm temperatures and mechanical stimuli(22, 23). We were unable to detect any differences in calcium transients elicited by the TRPV4 agonist GSK1016790A between Fabry and wildtype DRG neurons regardless of exposure to heat (**Supplemental Figure 2**).

### Heat stress does not increase Gb3 in Fabry DRG neurons

The heat shock response is known to cause increased ceramide synthesis(24, 25) and transient accumulation of glucosyl- and lactosyl-ceramide(26), precursors for Gb3, in nonneuronal cells. As Gb3 accumulates in Fabry disease and is an α-Gal A substrate, we asked whether neuronal and behavioral sensitization following heat exposure in Fabry disease was due to increased Gb3 load. As expected, Fabry DRG neurons had significantly more Gb3 than wildtype neurons (**Supplemental Figure 3**). Heat exposure did not increase Gb3 in either Fabry or wildtype neurons, however, suggesting that elevated Gb3 accumulation is not a major driver of increased TRPA1 sensitization or mechanical sensitization in Fabry rats exposed to heat.

### The heat shock response is impaired in Fabry DRG neurons

We next sought to understand why Fabry DRG neurons exhibited poor resilience to heat stress. The heat shock response is a network of molecular and metabolic pathways that contribute to cellular and organismal heat resilience(7, 8). Thus, we hypothesized that the heat shock response is impaired in Fabry DRG neurons. To test heat shock response induction, we assessed nuclear translocation of heat shock factor 1 (HSF1) in cultured DRG neurons (NeuN^+^) by confocal microscopy (**Figure 3**). At steady state, HSF1 is confined to the cytoplasm in both Fabry and wildtype DRG neurons. On exposure to heat in wildtype neurons, HSF1 translocates to the nucleus (Lamin B1^+^, **Figure 3A-D**), where it acts as the master transcriptional regulator of the heat shock response(9). Even following heat exposure, HSF1 is retained in the cytoplasm of Fabry DRG neurons, indicating impaired induction of the transcriptional phase of the heat shock response. Moreover, Fabry DRG neurons had lower expression of *Hspa1a* and *Hsp90aa1* (**Figure 3E-F**), the inducible paralogs of cytoplasmic HSP70 and HSP90. Together, these data indicate an impaired heat shock response in the peripheral nervous system of Fabry rats.

**Figure 3.**
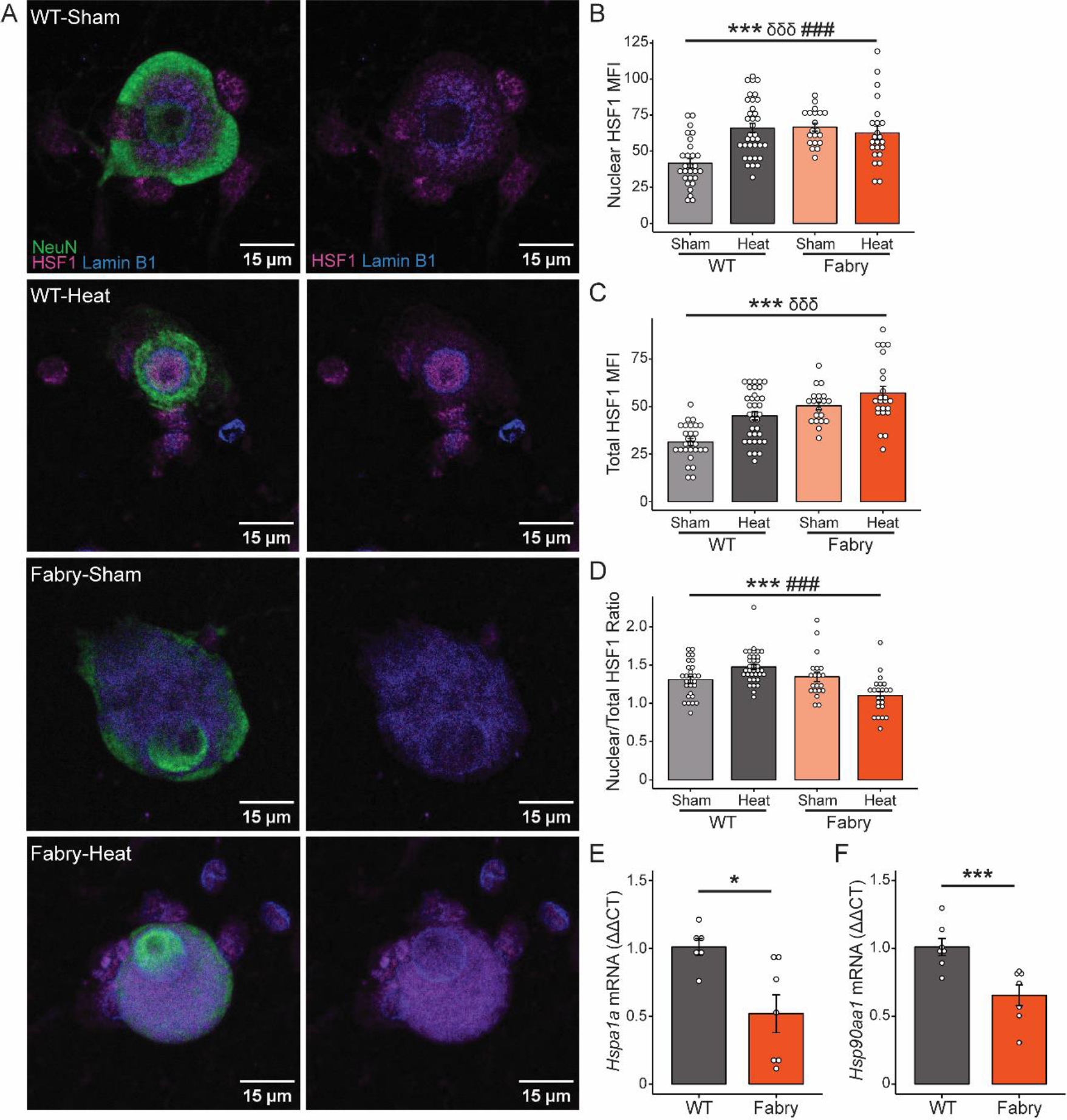
The heat shock response is impaired in Fabry DRG neurons. (A) Representative images of wildtype and Fabry DRG neurons (NeuN^+^) stained for translocation of heat shock factor 1 (HSF1) to the nucleus (Lamin B1^+^) following heat exposure. Mean fluorescence intensity (MFI) of HSF1 was measured in the nucleus (B) and whole soma (C) for each neuron and used to calculate a ratio between nuclear and total HSF1 (D) as a measure of HSF1 nuclear translocation (n=20-35 neurons cultured from 3 rats of each genotype). Fabry DRG express less *Hspa1a* (E) and *Hsp90aa1* (F) mRNA compared to wildtype DRG. (B-D) 2-way ANOVA, *** is genotype: p<0.005, δδδ is effect of heat: p<0.005, ### is genotype-heat interaction: p<0.005. (E-F) 2-tailed Student’s t-test, * p<0.05, *** p<0.005.

### HSP70 and HSP90 differentially regulate TRPA1 responses in naïve DRG neurons

As the heat shock response is known to confer neuroprotection and resilience to heat stress, we asked whether reduced heat shock response capacity could explain the sensitization of TRPA1 in Fabry DRG neurons. Naïve DRG neurons incubated overnight with a pan-HSP70 inhibitor (VER-155008) or a pan-HSP90 inhibitor (17-AAG) were exposed to transient heat prior to assessing calcium responses to AITC (**Figure 4A**). Incubation with VER-155008 (25μM) increased the proportion of AITC-responsive neurons at steady state and following heat stress (**Figure 4B-D**), indicating that HSP70 reduces TRPA1-mediated calcium transients. Conversely, overnight incubation with 17-AAG (0.5μM) decreased the percentage of AITC-responsive neurons regardless of heat exposure. These data may suggest that HSP90 facilitates TRPA1-mediated calcium transients; however, it is also possible that 17-AAG decreases the proportion of AITC-responsive DRG neurons through its ability to upregulate HSP70 and HSP90 expression(27, 28). To elucidate these mechanisms, we examined whether 30-minute exposure to VER-155008 or 17-AAG was sufficient to sensitize AITC-responses *in vitro* (**Figure 4E**). Indeed, acute incubation with either VER-155008 and 17-AAG increased the percentage of naïve DRG neurons that respond to AITC (**Figure 4G-H**), indicating that both HSP70 and HSP90 suppress TRPA1 activity.

**Figure 4.**
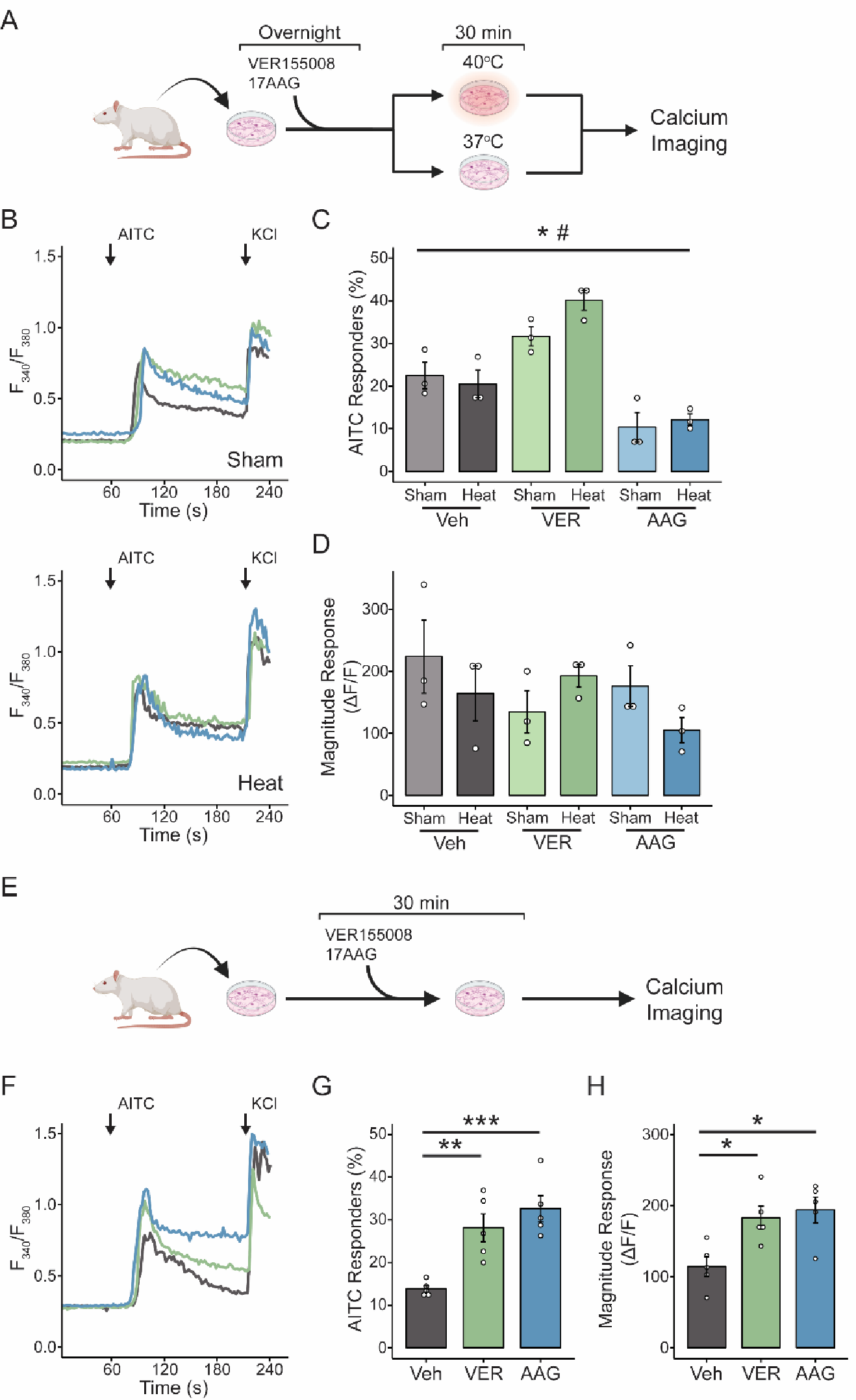
Heat shock proteins regulate TRPA1 activity. DRG neurons were cultured from Sprague-Dawley rats and incubated overnight in the HSP70 inhibitor VER15508 (VER) or the HSP90 inhibitor 17-AAG (AAG) prior to heat exposure and calcium imaging (A). (B) Representative traces of AITC-induced calcium transients from sham (*top*) and heat (*bottom*) treated DRG neurons. Incubation with VER increased the proportion of AITC-responsive neurons at baseline and following heat-treatment, while overnight AAG exposure decreased AITC responders regardless of heat (C). Neither VER nor AAG exposure overnight sensitized AITC-induced calcium transient magnitude (D). Each dot represents cultured DRG neurons per rat (n=3 rats, 50-90 neurons per rat). In a separate experiment, DRG neurons cultured from Sprague-Dawley rats were exposed to VER or AAG 30 minutes prior to calcium imaging (E). Representative traces are shown in (F). Acute exposure to both VER and AAG increased the proportion of AITC-responsive neurons (G) and the magnitude of AITC-evoked calcium transients (H) Each dot represents cultured DRG neurons per rat (n=4 rats, 50-90 neurons per rat). (C-D) 2-way ANOVA with repeated measures, * is effect of drug: p<0.05, # is effect of drug-heat interaction: p<0.05. (G-H) 1-Way ANOVA with repeated measures, p<0.05; Tukey’s *post hoc*, * is effect of drug: p<0.05, ** is effect of drug: p<0.01, ***is effect of drug: p<0.005.

### Heat shock protein-modifying treatments improve neuronal sensitization in Fabry disease

Having demonstrated that inhibiting the heat shock response is causes TRPA1 sensitization in naïve DRG neurons, we next asked whether improving the heat shock response in Fabry disease reduced TRPA1 sensitization. As overnight 17-AAG incubation increases expression of heat shock proteins(27, 28) and reduces AITC-responders (**Figure 4**), we incubated wildtype and Fabry DRG neurons with 17-AAG overnight prior to calcium imaging (**Figure 5A**). Overnight 17-AAG completely reversed the increased proportion of AITC-responders in Fabry DRG neurons at both steady state and following heat exposure (**Figure 5B-D**). Additionally, overnight incubation with recombinant human HSP70 (5 μg/mL) reduced both the proportion of AITC-responsive Fabry DRG neurons (**Figure 5F**) and the magnitude of AITC-induced calcium transients (**Figure 5G**), regardless of heat exposure. Thus, HSP70 and HSP90 suppress TRPA1 activity in both normal and pathological settings.

**Figure 5.**
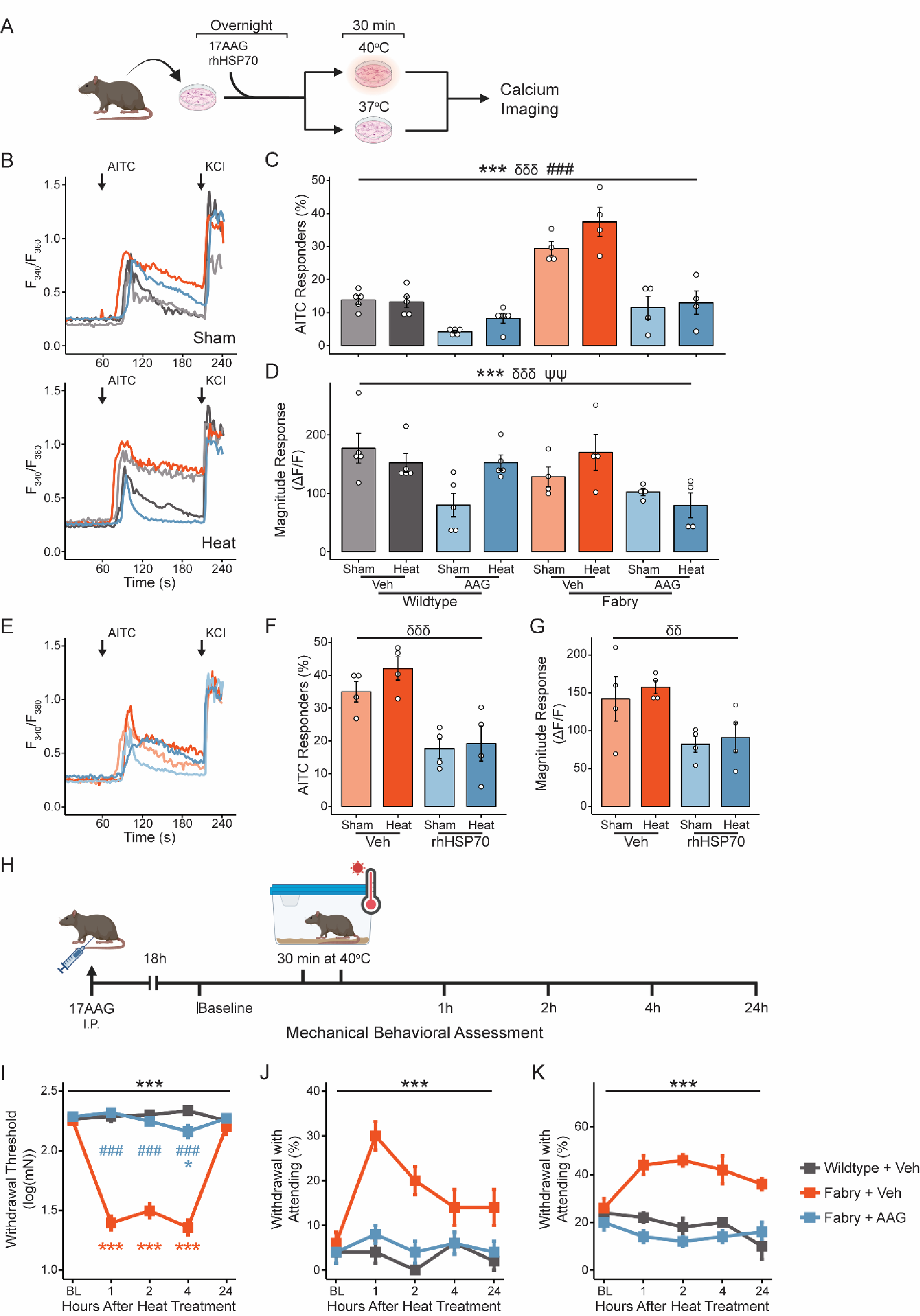
Targeting heat shock proteins reduces sensitization of TRPA1 and mechanical sensory behaviors in Fabry rats. DRG neurons cultured from wildtype or Fabry rats were cultured overnight in 17-AAG or recombinant human HSP70 (rhHSP70) prior to heat exposure and calcium imaging (A). (B) Representative traces of AITC-induced calcium transients from sham (*top*) and heat (*bottom*) treated DRG neurons exposed to AAG. Overnight AAG reduced the proportion of both wildtype and Fabry DRG neurons responsive to AITC regardless of heat exposure (C) and affected the magnitude of AITC-evoked calcium transients (D). Each dot represents cultured DRG neurons per rat (n=4-5 rats, 50-90 neurons per rat). Incubation with rhHSP70 also reduced the proportion of AITC-responsive Fabry DRG neurons (F) and the magnitude of AITC-evoked calcium transients (G) (Representative traces in (E)). Each dot represents cultured DRG neurons per rat (n=4 rats, 50-90 neurons per rat). Wildtype and Fabry rats were given an intraperitoneal injection of 17AAG 18 hours prior to mechanical behavioral testing before and after heat treatment (H). Heat exposure decreased mechanical withdrawal threshold (I) and increased aversive responses to dynamic brush (J) and noxious pinprick (K) in vehicle-injected, but not 17-AAG-injected, Fabry rats (n=5 per treatment group). (C-G) 3-Way ANOVA with repeated measures, * is genotype: p<0.05, *** is genotype: p<0.005, δδ is effect of drug: p<0.01, δδδ is effect of drug: p<0.005, ### is genotype-drug interaction: p<0.005, ψψ is genotype-drug-heat interaction: p<0.01. (I-K) 2-Way ANOVA with repeated measure, *** is group: p<0.005. (I) Tukey’s *post hoc*, *** is comparison to Wildtype-Veh: p<0.005.

### Heat shock protein-modifying treatments improve heat tolerance in Fabry rats

We next sought to determine whether altering the heat shock response with 17-AAG was able to prevent heat-evoked mechanical sensitization in Fabry rats. 12-week-old male hemizygous and female homozygous Fabry rats received a single intraperitoneal (IP) injection of 17-AAG (25 mg/kg) one day prior to mechanical behavioral testing (**Figure 5H**). At baseline, there were no statistically significant differences in mechanical withdrawal threshold or behavioral responses to noxious needle and dynamic brush stimuli (**Figure 5I-K**). Following heat treatment, vehicle-injected Fabry rats developed mechanical hypersensitivity compared to wild type animals as we had previously observed. However, prior injection with 17-AAG protected Fabry rats from developing hypersensitivity to any form of evoked mechanical stimulation, including von Frey filament testing and responses to dynamic brush and noxious needle stimulation. These data indicate that targeting HSP90 completely prevents environmental heat-induced mechanical hypersensitivity in Fabry rats. Thus, targeting HSP90 in Fabry disease may alleviate episodic pain in patients with Fabry disease.

## Discussion

Here we develop the first preclinical model of episodic pain in Fabry disease. Mild, transient environmental heat stress caused robust mechanical hypersensitivity in Fabry rats. We then used this model to explore molecular mechanisms underlying heat-evoked pain episodes in Fabry disease. Specifically, we determined that TRPA1, which is sensitized at steady state in Fabry disease, is further sensitized by heat in Fabry DRG neurons, while TRPV1 activity was unaltered by heat exposure. We next sought to determine the mechanisms causing heat to further sensitize TRPA1 in Fabry, but not wildtype, DRG neurons. We determined that the heat shock response was impaired in Fabry DRG, leading to lower expression of HSP70 and HSP90. Acute inhibition of HSP70 and HSP90 revealed that these molecular chaperones negatively regulate TRPA1 activity in naïve DRG neurons. Reciprocally, inhibition of HSP90 overnight, which is associated with altered fate for HSP90 client proteins(29, 30), completely prevented TRPA1 sensitization by heat in Fabry DRG neurons and heat-evoked mechanical hypersensitivity in Fabry rats.

Patients with Fabry disease often develop chronic pain in early adolescence and can experience heat-induced pain crises as early as the first two years of life(3, 4, 6), and carriers for Fabry disease also experience pain episodes(31, 32). Our prior work, however, demonstrated that baseline mechanical hypersensitivity develops in Fabry rats after 35 weeks of age, whereas heterozygous female Fabry animals never developed mechanical hypersensitivity(15). Here, we used young adult (*e.g.*, 10-14-week-old) Fabry rats to i) ensure that any hypersensitivity detected was due to heat-evoked sensitization, and ii) better model the early onset of heat-induced pain crises experienced by young patients (3–6). This model also recapitulates features of Fabry crises seen in patients. Fabry rats rapidly (less than one hour) developed mechanical hypersensitivity following brief exposure to moderate heat and, much like patient populations, these symptoms resolved within 24 hours. These features include hypersensitivity to presumably painful stimuli (noxious pinprick) as well as blunt and dynamic non-noxious mechanical stimuli (von Frey filaments and brush), which reflect the hyperalgesia and allodynia experienced by patients(4, 6), respectively. These results also imply that environmental variation may contribute to the discrepancy between onset of neuropathic pain in patients and pain behaviors in Fabry rats, which spend the majority of their lives sedentary and in climate-controlled housing. These factors may also contribute to the incomplete penetrance of chronic and episodic pain in Fabry carriers, as transient heat elicited similar sensitization in female heterozygous Fabry rats.

To the best of our knowledge, the work herein is among the first to use transient heat stress in rodents to model episodic pain. Environmental variation (*e.g.* exposure to heat) may also help explain discrepancies between the sensory presentations of other rodent models for episodic pain. Erythromelalgia is another inherited etiology of episodic pain in humans, during which heat exposure leads to burning, tingling pain in the affected body area(33, 34). Preclinical erythromelalgia is primarily studied in cellular expression systems(35–37) and the few mouse models that have been generated fail to show elevated pain-like behaviors relative to controls(38, 39). Importantly, these sensory behaviors were tested without heat stress, leaving open the possibility that heat would precipitate pain-like phenotypes akin to those experienced in clinical populations. Patients with sickle cell disease (SCD) also exhibit temperature-evoked painful episodes, and although episodic pain in SCD is associated more with colder temperatures, up to 26% of children with SCD report transitions from cold to warmth as a trigger for their pain(40). Simone and colleagues recently modeled episodic pain in SCD by exposing a mouse model of SCD to transient, mild cold stress(41, 42). These mice did not display pain-like behaviors at baseline, supporting the notion that ambient temperature variation can precipitate episodic pain. It is crucial we develop a better understanding of how ambient temperature affects the function of the sensory nervous system and provokes episodic pain, especially as the progression of climate change and global temperature variation with warming(43) will increase the frequency of heat-evoked pain episodes.

Although we previously defined a role for TRPA1 in chronic pain associated with Fabry disease(15), our current findings are the first to demonstrate a role for TRPA1 as a contributor to episodic pain in Fabry disease. TRPA1 has been implicated in at least two other episodic pain conditions. Kremeyer, *et. al*. described a TRPA1 gain-of-function mutation in a Columbian family with Familial Episodic Pain Syndrome(44). Episodic pain resulting from this mutation was not triggered by heat, but by cold exposure, exercise, and illness. Additionally, we have hypothesized previously that TRPA1 could contribute to episodic pain in SCD through sensitization by reactive oxygen and nitrogen species, lipid mediators, and hypoxia(45). Indeed, single nucleotide polymorphisms in *TRPA1* (rs920829) are associated with acute pain in SCD(46). Thus, establishing heightened sensitivity of TRPA1 in Fabry DRG neurons following heat stress continues to build evidence toward a model in with TRPA1 integrates cell stress and contributes to episodic pain.

Pain, when not maladaptive, is instructive for the avoidance of noxious stimuli and environmental stressors. It is therefore logical that molecular networks responsible for conferring resilience to stress, such as the heat shock response, also regulate sensory neuronal activity. Others have shown that HSP70 family members regulate the trafficking and function of TRPV1(14) and acid-sensing ion channel 2(47). Notably, Iftinca, *et. al.* did not report inhibition of TRPA1 by heat stress alone in a cell expression system(14), consistent with our observations here. Rather, we demonstrate that inhibition of HSP70 disinhibits TRPA1, which is exacerbated by heat. Moreover, we demonstrate a time-dependent regulation of TRPA1 by HSP90 inhibition. Acute inhibition of HSP90 by 17-AAG lead to sensitization of TRPA1, which may be associated with loss of refoldase activities of HSP90(29, 48, 49) and akin to the effect of HSP70 inhibition. Longer exposure to 17-AAG is associated with a shift in chaperone-cochaperone complexes, however, and is associated with polyubiquitination and degradation(29, 30). It is therefore possible that pretreatment with 17-AAG affects the trafficking and degradation of TRPA1 or known facilitators of TRPA1, such as protein kinase A, protein kinase C(50–52), or mitogen-activated proteins kinases(53, 54). Together, our findings identify HSP70 and HSP90 as a previously undefined regulatory axis for TRPA1. This finding is of critical importance to the somatosensory field given the role of TRPA1 not only in pain associated with Fabry disease(15), but also with itch and many other inflammatory and neuropathic pain conditions(18). Thus, more work is needed to define exactly how modulating the activities of heat shock proteins regulates the activity of TRPA1 and other TRP channels.

Another question that remains is the extent to which the heat shock response is altered in Fabry disease. We demonstrated impaired initiation of the heat shock response, as HSF1 was unable to translocate to the nucleus of Fabry DRG neurons following heat exposure. This deficit is congruent with the decreased *Hspa1a* and *Hsp90aa1* transcripts we observed in Fabry DRG, as HSF1 is the master transcriptional regulator of the heat shock response(7, 9). It is also possible the heat shock response is dysregulated in Fabry disease beyond the expression level.

Nikolaenko *et. al.* demonstrated direct interaction between lyso-Gb3, the product of Gb3 deacylation, and multiple heat shock proteins, including HSP70 and HSP90 paralogs(55). It is possible this interaction leads to altered localization of heat shock proteins or affects their foldase and holdase activities, especially near the membrane given the amphiphilic nature of glycosphingolipids; however, the functional consequences of Gb3/lyso-Gb3 interaction with heat shock proteins remains to be explored.

Heat shock proteins regulate pain beyond simply regulation of ion channels. Streicher and colleagues have demonstrated an HSP90-isoform specific regulation of antinociception conferred by opioids(56, 57), but not gabapentinoids(56), in the spinal cord. Thus, it is possible that an impaired heat shock response in Fabry rats also affects descending inhibitory and facilitatory spinal pain pathways, though no studies to date have examined spinal pain processing in Fabry disease. HSP70 and HSP90 also act as damage-associated molecular patterns when released by damaged cells(58, 59) and facilitate toll-like receptor 4 (TLR4) signaling(60, 61). These signaling events are thought to be pronociceptive(61, 62). Indeed, exogenous HSP90 evokes calcium transients when acutely applied to cultured DRG neurons(63), though it is unclear whether this effect is mediated through TLR4 engagement. This is particularly notable in Fabry disease, as elevated TLR4 signaling occurs as a result of Gb3 accumulation(64), and elevated tumor necrosis factor α (TNFα), a downstream result of TLR4 signaling(65), is elevated in the sera of patients with Fabry disease following a pain crisis. Though heat shock is known to increase ceramide and glycosphingolipid accumulation(24–26), this is unlikely to be a direct driver of elevated TLR4 and TNFα signaling as we did not observe increased Gb3 accumulation in Fabry DRG following heat stress.

This study establishes the first preclinical model for episodic pain in Fabry disease, allowing for the first time meticulous dissection of the molecular mechanisms underlying this debilitating lifelong condition. We determined that the heat shock response is impaired in Fabry DRG sensory neurons, which lead to the sensitization of TRPA1 by heat and, in turn, mechanical hypersensitivity. In doing so, we discovered that HSP70 and HSP90 are potent regulators of TRPA1. Strikingly, we found that inhibition of HSP90 completely prevented both neuronal sensitization and all forms of mechanical hypersensitivity evoked by heat in Fabry disease. Therefore, these findings not only identify a novel, therapeutically viable regulatory mechanism for TRPA1, but they demonstrate the incredible efficacy of targeting the heat shock response as a therapeutic strategy for episodic pain in Fabry disease.

## Methods

### Sex as a Biological Variable

Fabry disease is an X-linked genetic disorder that primarily affects males. Despite this, female patients with homozygous loss-of-function mutations in *GLA* and heterozygote “carriers” do experience neurological complications, including heat-induced pain crises. As such, pain behaviors were assessed in all five possible genotypes (male: *Gla^+/0^*, *Gla^−/0^*; female: *Gla^+/+^*, *Gla^+/−^*, *Gla^−/−^*). Both male and female Sprague-Dawley outbred rats were used throughout experimentation.

### Animal Model and Institutional Approval

The X-linked genetic Fabry disease rat model (Rat Genome Database symbol: GLAem2Mcwi) and age-matched wildtype littermate controls were used for sensory behavioral experiments. These animals also provided primary sensory neurons for calcium imaging and immunofluorescent experiments. Both male and female rats were used between 12 and 35 weeks of age. Sprague-Dawley outbred rats (Taconic) aged 12-30 weeks of age provided primary sensory neurons for calcium imaging experiments examining the role of heat shock proteins on TRPA1 activity (naïve rats). All animals were maintained on a 12:12 light:dark cycle in the Medical College of Wisconsin (MCW) Biomedical Resource Center, and all experiments involving live animals were approved by the MCW Institutional Animal Care and Use Committee (#5188).

### Transient Heat Treatment

For *in vivo* application of transient heat treatment, rats were placed in a rat cage modified by the MCW Biomedical Engineering Department to include a small metal grate protecting a heating element and fan. This heat box was also fitted with a thermometer and temperature control (InkBird), allowing for stable temperature maintenance at 40°C for 30 minutes. No more than three rats were heat-treated in the same heat box at the same time, and only cage-mates received heat treatment together. For *in vitro* application of transient heat treatment, cultured dorsal root ganglia (DRG) neurons were place in a cell culture incubator set to 40°C for 30 minutes. For intraperitoneal (I.P.) 17-AAG injection, a 10 mM stock of 17-AAG (MedChemExpress) or DMSO was diluted in 0.9% sterile saline to a working concentration of 0.213 mM (125 mg/L). Rats were weighed, briefly anesthetized under isoflurane, and injected with 25 mg/kg 17-AAG.

### Sensory Behavioral Testing

All rodent sensory behaviors were assessed in the MCW Neuroscience Resource Center rat behavior suite. Rat plantar cutaneous mechanical sensitivity was assessed as previously described. Briefly, rats were acclimated to the behavioral facilities and behavioral testing apparatuses (small acrylic single-housing chambers atop a wire mesh table) for at least 1 hour each at least twice prior to sensory testing in the presence of white noise. Mechanical withdrawal threshold was assessed by the “up-down” method using calibrated von Frey filaments. Response to noxious pinprick (needle) and dynamic brush (paintbrush) were then assessed as previously described(15, 66). Briefly, stimuli were applied to the plantar surface of the right and left hind paws 5 times each. The response to each individual stimuli were classified in three ways: 1) null response, 2) simple withdrawal and return of the stimulated paw, and 3) paw withdrawal with attending or aversive response (paw flutter, shaking, hovering, guarding, licking, etc.). All behavioral experiments were performed by a blinded experimenter, who remained blinded until the conclusion of behavioral testing and analysis of resultant data.

### Isolation and Culture of Primary Dorsal Root Ganglion (DRG) Neurons

DRG neurons were harvested, dissociated, and cultured as previously described(67). Briefly, lumbar DRG neurons were acutely dissected and cleaned of neurites and the epineurium immediately following euthanasia by isoflurane inhalation and decapitation with guillotine. Whole DRG were then incubated in collagenase IV (1 mg/mL) in Dulbecco’s Minimum Essential Media (DMEM) for 45 minutes at 37°C. DRG were then incubated in trypsin (0.05%) for 45 minutes at 37°C, which was immediately neutralized with horse serum, washed in DMEM, and resuspended in complete DRG media (DMEM/F12, 10% fetal horse serum, 1% glucose, 2 mM L-glutamine, 100 U/mL penicillin, and 100 μg/mL streptomycin). DRG were progressively mechanically dissociated with a p1000 and p200 micropipette prior to plating on laminin-coated glass coverslips. Between 1-2 hours after plating, dissociated DRG neurons were fed with complete DRG media or complete DRG media supplemented with VER155008 (25 μM, MedChemExpress), 17-AAG (0.5 μM, MedChemExpress), or recombinant human HSP70 (5 μg/mL, R&D Systems, lot #DBPG0323101).

### Calcium Imaging

Dissociated DRG neuronal soma were used for calcium imaging 18-24 hours after plating. Somata were washed with extracellular buffer (in mM: 150 NaCl, 10 HEPES, 8 glucose, 5.6 KCl, 2 CaCl_2_, 1 MgCl_2_; pH 7.40 +/− 0.03, mOsm 320 +/− 3) for 1-2 minutes, incubated in 2.5 mg/mL Fura-2 (Life Technologies) in 2% bovine serum albumin for 45 minutes, and washed with extracellular buffer for 30 minutes. For acute exposure to heat shock protein inhibitors, this wash step was performed in extracellular buffer containing VER155008 (25 μM) or 17-AAG (0.5 μM). Fura-2 fluorescence was measured with excitation at 340 nm and 380 nm and captured by a cooled Andor Zyla-SCMOS cameral (Oxford Instruments). NIS Elements software (Nikon) was used to analyze 340 and 380 signals and calculate bound/unbound calcium ratio (340/380). To induce intracellular calcium transients, DRG soma were exposed to allyl isothiocyanate (AITC, 30 μM, 30 seconds), GSK1016790A (GSK101, 30 nM, 30 seconds), or capsaicin (50 nM, 30 seconds) followed by 2-minute washout with extracellular buffer. Cell health was assessed by transient (30 second) exposure to 50 mM KCl. Soma that exhibited at least 20% increase in 340/380 ratio over baseline were considered positive responders. Cells were only included for analysis if they were responders to either the experimental stimulus or 50 mM KCl.

### Immunofluorescent Microscopy

Immunofluorescent microscopy *in vitro* with dissociated cells was performed using modifications from a previously described protocol(67, 68). Dissociated cells were adhered to glass coverslips overnight. Prior to imaging, coverslips were washed in filter phosphate buffered solution (PBS), fixed by 4% paraformaldehyde for 20 minutes at room temperature, and subject to three 5-minute washes with PBS. Coverslips were then incubated in blocking solution comprised of SuperBlock, 1.5% normal donkey serum, and 0.5% Triton X-100 for 1-2 hours at room temperature. After blocking, coverslips were incubated with primary antibodies (**Supplemental Table 1**) diluted in SuperBlock and blocking solution (100 μL Superblock to 100 μL blocking solution) overnight at 4°C. The next morning, coverslips were washed three times for 5 minutes in filtered PBS. Coverslips were then incubated in secondary antibodies (**Supplemental Table 1**) diluted in SuperBlock for 1 hour protected from light. After three additional washes with filtered PBS, coverslips were then mounted to SuperFrost Plus glass slides with ProLong Gold and sealed with nail polish and superglue. Slides were allowed to cure for at least 24 hours prior to imaging.

Microscopy images were acquired using an A1R Nikon confocal microscope equipped with 405 nm, 488 nm, 560 nm, and 640 nm laser lines using 20x and 40x air objectives. Z-stacks (8-12 μm) were taken for images of dissociated cells and tissue sections, and maximum intensity projections were generated from z-stacks using NIS Elements for downstream data analysis. Maximum intensity projections were analyzed using the FIJI version of ImageJ by a blinded observer. For analysis of Gb3 accumulation, regions of interest (ROIs) were hand-drawn around neuronal soma marked by NeuN and mean fluorescence intensity (MFI) was collected. For analysis of HSF1 nuclear translocation, ROIs were drawn around whole neurons (NeuN^+^) and neuronal nuclei (encircled by lamin B1). MFI were collected for both whole neurons and neuronal nuclei. The nuclear HSF1 ratio was calculated by dividing nuclear HSF1 MFI and total HSF1 MFI for each individual cell. Representative images were generated using a standard deviation projection of z-stacks with background subtraction in across all channels. All analyses were performed by a blinded experimenter, who remained blinded until completion of image processing and data analysis.

### Reverse Transcriptase Quantitative Polymerase Chain Reaction (RT-qPCR)

RNA was isolated from L3-6 DRG harvested, dissociated, and cultured according to the above methods. 24 hours post-plating, DRG cultures were washed 3 times with ice-cold sterile PBS and lysed in TRIzol for 5 minutes at room temperature. RNA was isolated from the resultant lysate using PureLink Mini Prep kits (ThermoFisher) according to manufacturer’s instructions and quantified by NanoDrop. RNA was then converted into copy DNA (cDNA) using 5X VILO reaction mix and 10X SuperScript Reverse Transcriptase enzyme mix. Resultant cDNA was then diluted in water and mixed with the appropriate primers (**Supplementary Table 2**) and SYBR Green MasterMix for quantitative PCR in a 384-well format. Cycle thresholds were converted to ΔΔCT using *B2m* as a loading control.

### Statistical Analyses

All statistical analyses were performed using the R programming language (version 4.3.2) and the RStudio environment. Longitudinal experiments were analyzed by 2-way mixed-models analysis of variance (ANOVA) with repeated measures. Data were otherwise analyzed by *N*-way ANOVA, where *N* is the number of independent variables of the experiment. Significant main and interaction terms were further analyzed using Tukey’s honest significant difference *post hoc* test. Confidence intervals are presented as standard error of the mean.

## Acknowledgments

We would like to thank Bonnie Freudinger and the Medical College of Wisconsin (MCW) Engineering core the design and fabrication of the rat heating apparatus used in this manuscript. We would like to thank the MCW Neuroscience Resource Center for behavioral suite access, and the MCW Cardiovascular Research Center for access to the Nikon A1R confocal microscope. We thank Dianise Rodríguez García for her helpful comments on figure design. We would like to thank Drs. Katelyn Sadler, Tayler Sheahan, Vanessa Ehlers, and Brian Lin for their helpful insight during initial editing for this manuscript. This work was funded by National Institutes of Health grants R37NS108278 (CLS), R01NS070711 (CLS), and F32NS138223 (JDE), and support from the Advancing a Healthier Wisconsin Endowment (JDE and CLS).

## Contributions

JDE and CLS conceptualized the experiments. JDE and EKP developed the methodology. JDE, EKP, SMP, and AS conducted the experiments. JDE analyzed the data. JDE created the original manuscript draft and figure designs. All authors contributed to editing this paper. Supervision was provided by CLS. Funding for the work was acquired by JDE and CLS.

## Data Availability

By request.

