## Supplemental Figure 1 for "Episodic pain in Fabry disease is mediated by a heat shock protein-TRPA1 axis"

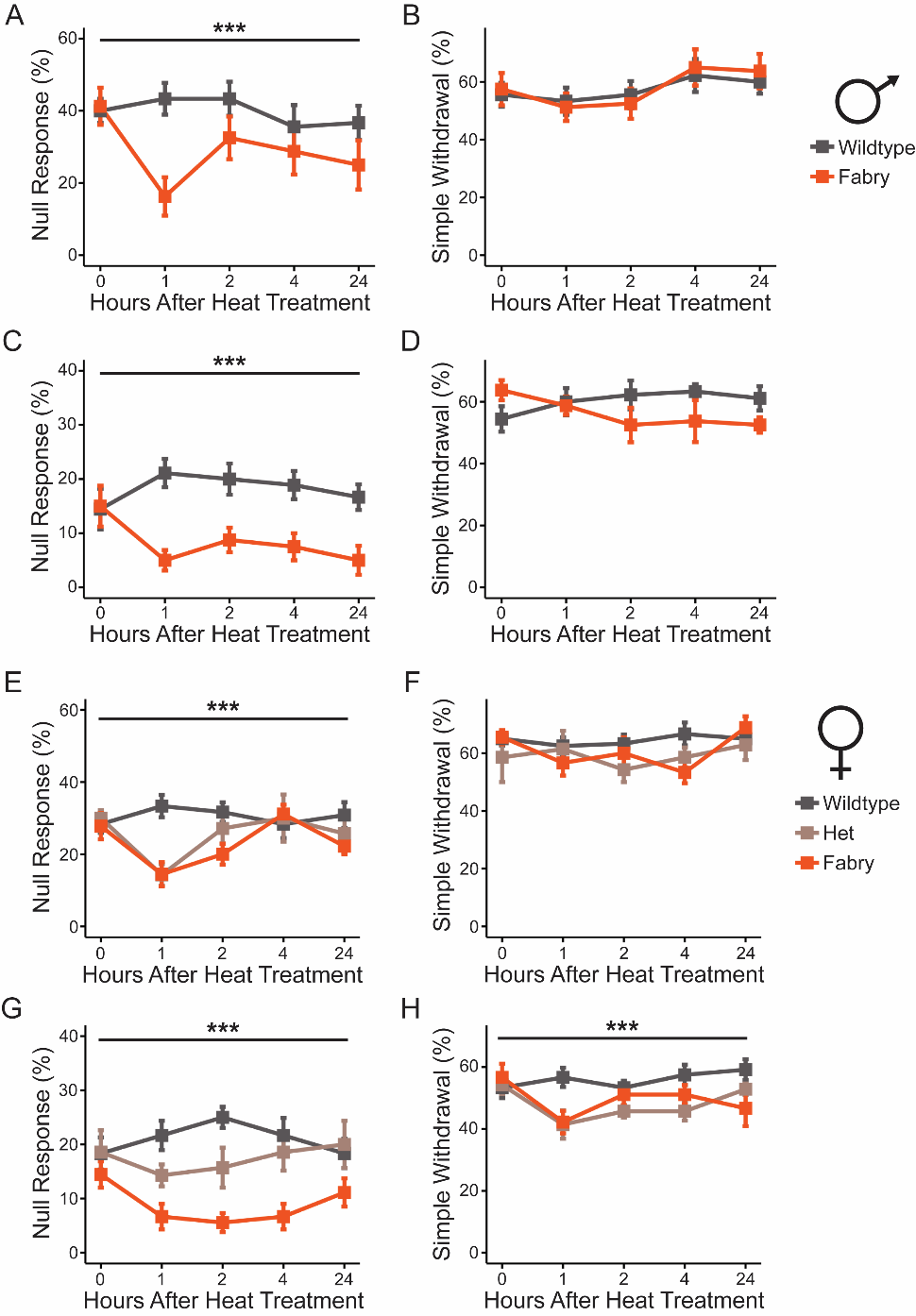

**Supplemental Figure 1. Heat decreases null responses to brush and needle in Fabry rats.** Male Fabry rats exhibited fewer null responses (A), but no change in simple withdrawal (B), to brush following heat exposure. They similarly demonstrated fewer null responses to noxious pinprick (C) with no change to simple withdrawal (D). Heterozygous and homozygous female Fabry rats demonstrated similar responses to brush (E-F) and needle (G-H), though there were significant reductions in simple withdrawal to needle in heated Fabry female rats (H). 2-Way ANOVA with repeated measures, *** indicates effect of genotype: p < 0.005.
