## Supplemental Figure 2 for "Episodic pain in Fabry disease is mediated by a heat shock protein-TRPA1 axis"

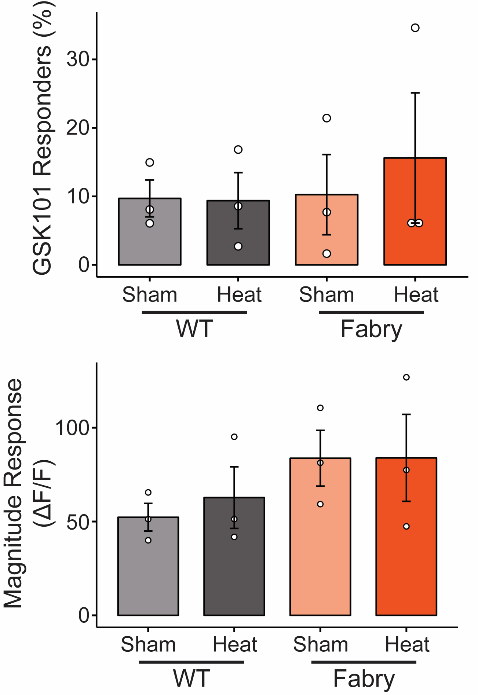

**Supplemental Figure 2. Heat exposure does not sensitize TRPV4 in Fabry DRG.** Neither the percent responders to TRPV4 agonist GSK1016790A (*top*) nor the magnitude of calcium flux (*bottom*) were different between wildtype and Fabry DRG neurons with or without heat treatment. 2-way ANOVA. Each dot represents cultured DRG neurons per rat (n=3 rats, 50-90 neurons per rat).
