## Supplemental Figure 3 for "Episodic pain in Fabry disease is mediated by a heat shock protein-TRPA1 axis"

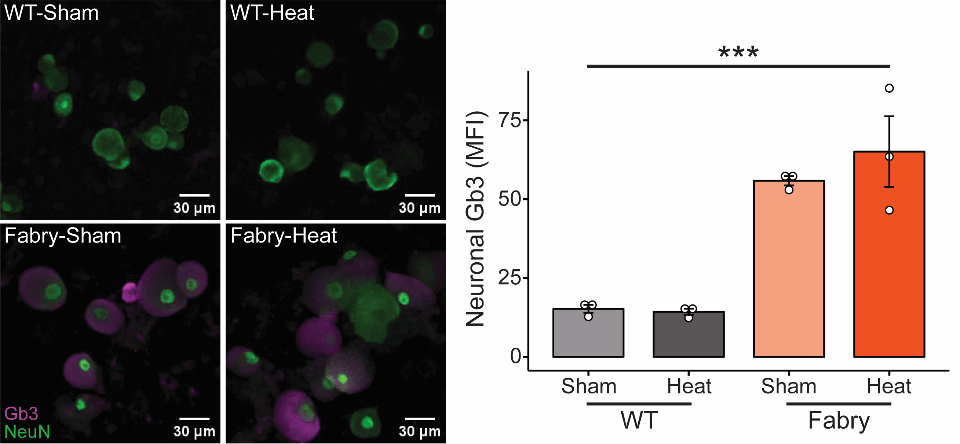

**Supplemental Figure 3. Heat exposure does not increase globotriaosylceramide accumulation in Fabry DRG neurons.** (*left*) Representative images of DRG neurons (NeuN^+^) cultured from wildtype and Fabry rats stained for CD77/globotriaosylceramide (Gb3) following heat treatment. Quantification of neuronal Gb3 by mean fluorescence intensity (*left*, MFI) demonstrates that increased Gb3 in Fabry DRG neurons is not exacerbated by heat. 2-way ANOVA, *** is genotype: p<0.005. Each dot represents cultured DRG neurons per rat (n=3 rats, 50-90 neurons per rat).
