## Supplemental Table 1 for "Episodic pain in Fabry disease is mediated by a heat shock protein-TRPA1 axis"

| **Supplemental Table 1: Antibodies used in these studies.** | | |
| --- | --- | --- |
| *Primary Antibodies* | | |
| NeuN | Sigma-Aldrich (ABN90) | 1:1000 |
| HSF1 | Cell Signaling Technologies (#4356) | 1:200 |
| Lamin B1 | ProteinTech (66095) | 1:300 |
| CD77/Gb3 | FisherScientific (50-204-1196) | 1:250 |
| *Secondary Antibodies* | | |
| Donkey α-Rabbit AF647 | Invitrogen (A-31573) | 1:1000 |
| Donkey α-Mouse AF405 | Invitrogen (A48257) | 1:1000 |
| Donkey α-Guinea Pig AF488 | Jackson ImmunoResearch (AB_2340472) | 1:1000 |
