## Supplemental Table 2 for "Episodic pain in Fabry disease is mediated by a heat shock protein-TRPA1 axis"

| **Supplemental Table 2: Primers used in these studies.** | |
| --- | --- |
| *B2m* – F | 5’-AATTCACACCCACCGAGACC-3’ |
| *B2m* – R | 5’-TGATTACATGTCTCGGTCCCA-3’ |
| *Hspa1a* – F | 5’-ATCGAGGAGGTGGATTAGAG-3’ |
| *Hspa1a* – R | 5’-ACCGAACGAAGGAGTTAATG-3’ |
| *Hsp90aa1* – F | 5’-GCTTGACCGACCCTAGTAAA-3’ |
| *Hsp90aa1* – R | 5’-CCAATGCCAGTATCCACAATAG-3’ |
